# Structural and enzymatic differences between human MDH isoforms underlie differential sensitivity to metabolic intermediates

**DOI:** 10.1101/2025.07.15.665002

**Authors:** Chris Berndsen, Angela J. Kayll, Ruhi Rahman, Jackson R Tester, Trip Beaver, Caroline Tuck, Sonja Neve-Hoversten, Lisa Gentile, Tamerat Mandanis, Joseph J Provost

## Abstract

Malate dehydrogenase (MDH) catalyzes the reversible redox reaction of malate and oxaloacetate, linking cellular redox balance to central metabolism. In mammals, this reaction is carried out by cytosolic and mitochondrial MDH, which share a conserved catalytic mechanism but function in distinct metabolic environments. Although structural and kinetic differences between these isoforms have been described, their relative sensitivity to endogenous metabolic intermediates remains incompletely understood. Here, we compared recombinant human cytosolic MDH (hMDH1) and recombinant human mitochondrial MDH (hMDH2) under matched experimental conditions to evaluate the effects of NAD^+^, ADP, ATP, citrate, α-ketoglutarate, (αKG) and glutamate. Enzymatic assays revealed that hMDH2 is inhibited by a broader range of metabolites than hMDH1, with particularly pronounced differences in sensitivity to ATP, citrate, and NAD^+^. Fluorescence measurements showed that only a subset of these metabolites produced detectable changes in the local environment of the fluorescence reporters, indicating that functional inhibition does not necessarily correspond to a measurable fluorescence response. Small-angle X-ray scattering (SAXS) demonstrated that both isoforms remain dimeric in solution without detectable large-scale structural rearrangements upon ligand addition. Structural comparisons and computational analyses suggest that differences in local conformational dynamics and active-site environments may contribute to their distinct responses. Together, these findings establish that human MDH1 and MDH2 differ in their sensitivity to metabolic intermediates despite their conserved catalytic architecture. This isoform-specific sensitivity may contribute to differences in how cytosolic and mitochondrial MDH activities respond to changes in cellular metabolic conditions.

## 1. Introduction

Malate dehydrogenase (MDH) is an essential, conserved enzyme that catalyzes the reversible interconversion of malate and oxaloacetate, coupled to NAD^+^/NADH redox cycling (Bernstein LH, and Everse J. 1978, De Lorenzo L et al. 2024, Goward CR, and Nicholls DJ. 1994 and McCue W.M. and Finzel B.C., 2022). In mammalian systems, this reaction is carried out by two primary isoforms: cytosolic MDH1 and mitochondrial MDH2. Although these isoforms share a conserved catalytic mechanism and overall structural features, they operate in distinct metabolic environments, supporting different aspects of cellular metabolism.

Human mitochondrial MDH (hMDH2) functions in the mitochondrial matrix as part of the tricarboxylic acid (TCA) cycle, where it contributes to NADH production for oxidative phosphorylation. In contrast, human cytosolic MDH (hMDH1) operates in the cytosol, where it supports the malate–aspartate shuttle and contributes to redox balance and biosynthetic processes. These roles position the two isoforms at key junctions of energy metabolism and suggest that they may respond differently to changes in metabolite availability and redox state.

Despite their shared chemistry and general structural fold, hMDH1 and hMDH2 differ substantially in primary sequence and exhibit structural differences that may contribute to their distinct biochemical properties. Both isoforms function as homodimers and share the conserved two-domain dehydrogenase architecture, consisting of an N-terminal Rossmann-fold nucleotide-binding domain and a C-terminal region involved in substrate binding and catalysis (Figure 1A and B). Sequence alignment and structural comparison of hMDH1 (PDB: 7RM9) and hMDH2 (PDB: 2DFD) reveal that regions of sequence divergence can retain similar backbone conformations, whereas other regions exhibit substantial structural displacement.

**Figure 1.**
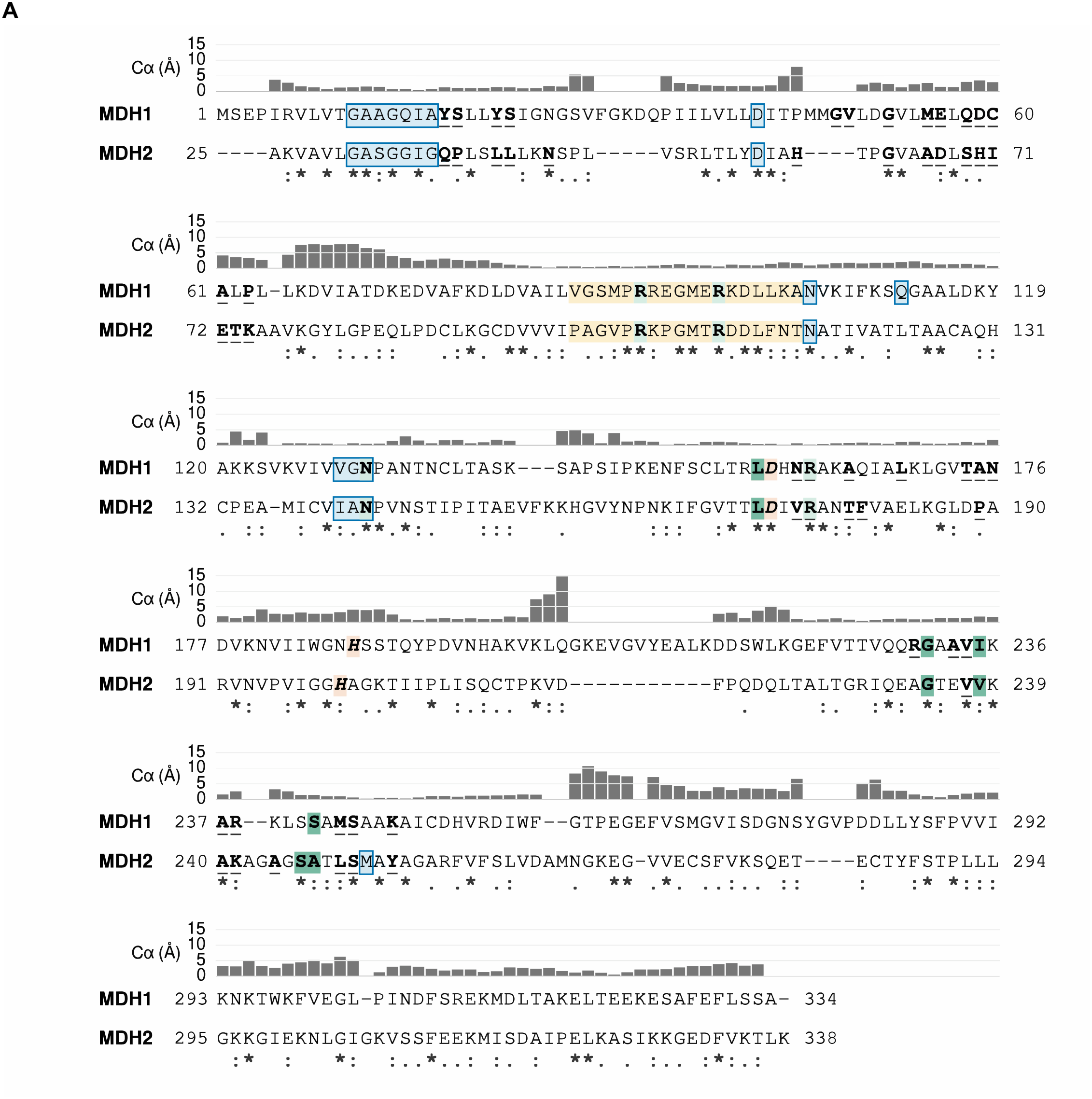
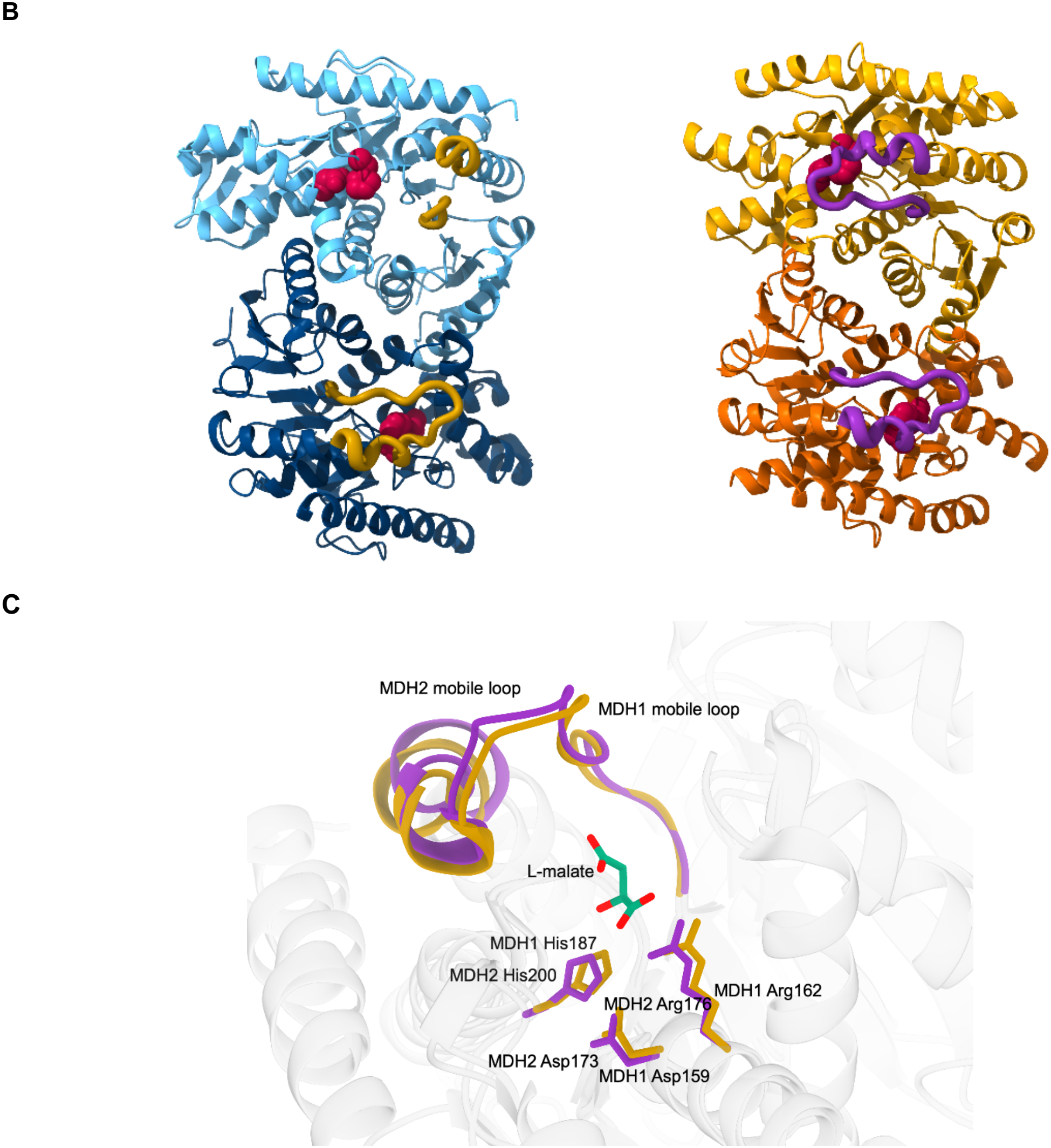
Sequence and structural correspondence of human MDH1 and MDH2. **(A) Sequence alignment and structural comparison of hMDH1 and hMDH2.**The UCSF ChimeraX Matchmaker alignment compares chain A of hMDH1 (PDB: 7RM9) and hMDH2 (PDB: 2DFD). hMDH1 uses canonical residue numbering (UniProt P40925-1, residues 1–334), whereas hMDH2 uses precursor numbering (UniProt P40926-1, residues 25–338), excluding the mitochondrial targeting peptide and expression tag. Hyphens indicate alignment gaps. Consensus symbols indicate identical (*), strongly similar (:), and weakly similar (.) residues based on Clustal amino acid groups. Gray bars indicate distances between corresponding Cα atoms following a single global structural superposition, using a consistent 0–15 Å scale across all alignment blocks. Columns without paired Cα atoms are left blank. These distances represent structural differences between the aligned crystal structures, not local refitting or conformational fluctuations. Light blue shading and blue outlines identify NAD-binding residues, with consecutive contacts enclosed in a single box. Pale yellow shading identifies the approximate mobile active-site loops (hMDH1 residues 87–105; hMDH2 residues 99–117). Pale green shading with bold lettering identifies substrate-binding residues, while darker green shading identifies additional residues within 4 Å of the bound substrate or analogue. Salmon shading with bold italic lettering identifies the catalytic Asp/His pairs (hMDH1 D159/H187; MDH2 D173/H200). Bold lettering with gray underlines identifies residues at the A–B dimer interface, defined using a minimum buried surface area of 15 Å^2^ per residue across both subunits. Ligand contacts were determined from minimum heavy-atom distances to malonate in hMDH1 (7RM9, chain A) and L-malate in hMDH2 (2DFD, chains A and B). hMDH2 residue numbers were converted to precursor numbering by adding 18 to the residue numbers. Where annotations overlap, NAD-binding residues retain blue outlines, and ligand-binding or catalytic residue shading takes precedence over mobile-loop shading. Residue proximity to a ligand does not necessarily establish a functional role in binding. **(B) Dimer architecture of hMDH1 and hMDH2**. Cartoon representations of hMDH1 (7RM9, left) and the A–B dimer of hMDH2 (2DFD, right), shown in matched orientations following structural superposition. Subunits are distinguished by light and dark blue for hMDH1 and gold and orange for MDH2. The mobile-loop regions are highlighted in gold (hMDH1, residues 87–105) and purple (hMDH2, residues 99–117). Catalytic Asp and His side chains are shown as crimson spheres: Asp159 and His187 in hMDH1, and Asp173 and His200 in hMDH2. hMDH2 residue numbers refer to the full-length precursor sequence. **(C) Structural comparison of the hMDH1 and hMDH2 active sites**. Superposition of chain A from hMDH1 and hMDH2. Mobile-loop regions and selected active-site residues are colored gold for hMDH1 and purple for hMDH2; the surrounding backbone is shown in translucent gray. Catalytic Asp and His residues and the substrate-binding Arg residue are displayed as sticks and labeled. Malonate from 7RM9 is omitted. hMDH2 labels use full-length precursor numbering. The overlay illustrates the corresponding active-site arrangements and mobile-loop positions in these crystal structures.

The N-terminal glycine-rich nucleotide-binding motif and residues comprising the catalytic machinery exhibit relatively close structural correspondence. In contrast, several extended regions within both the nucleotide-binding and catalytic domains, including regions corresponding to hMDH1 residues 66–76 and 260–277, exhibit Cα displacements exceeding 5 Å at multiple positions following global structural superposition (Figure 1A). Overlay of the active sites further illustrates conservation of the catalytic residues alongside differences in the sequence and local structural organization of the surrounding regions (Figure 1C). Such differences may influence conformational dynamics, substrate recognition, and communication between subunits without altering the fundamental catalytic architecture. Considered alongside their distinct metabolic functions and compartment-specific biochemical environments, these differences provide a structural and physiological rationale for investigating isoform-specific regulation by endogenous metabolic intermediates

Of particular interest in this context is the “thumb” loop, a structural element adjacent to the mobile active-site loop that spans the dimer interface and has been proposed to coordinate conformational changes between monomers. Flexibility in this region may facilitate communication between active sites, providing a potential mechanism for allosteric regulation and modulation of enzyme activity in response to shifting metabolic demands. Differences in the sequence and structural organization of the thumb and adjacent mobile active-site loop may therefore contribute to isoform-specific regulatory properties. Although this model has not been established experimentally for human MDH1 and MDH2, it highlights the potential importance of local conformational dynamics and intersubunit communication in enzyme regulation.

Kinetic differences between the isoforms are well documented and align with their distinct metabolic functions. Both hMDH1 and hMDH2 follow an ordered bi-bi reaction mechanism in which NAD^+^ binds prior to malate and NADH is released as the final product. Reported kinetic parameters indicate that hMDH1 generally exhibits higher apparent affinity for oxaloacetate, whereas hMDH2 supports higher catalytic turnover, consistent with their respective cytosolic and mitochondrial roles (Fahien L.A. et al. 1988; Lorenzo L. et al. 2024; Martinez-Vas B.M. et al. 2024). These differences are typically reflected in lower micromolar K_m_ values for oxaloacetate in hMDH1 relative to hMDH2 and modestly higher turnover rates for hMDH2, supporting sustained flux through the tricarboxylic acid cycle. Together, these features suggest adaptive specialization of each isoform to its metabolic environment. However, much of the available kinetic and regulatory data derives from non-human systems or from studies performed under variable experimental conditions, including differences in pH, substrate concentrations, and reaction direction, making direct comparison between human MDH1 and MDH2 difficult. As a result, comprehensive analyses using recombinant human proteins under matched biochemical and structural conditions remain limited.

While structural and kinetic differences between hMDH1 and hMDH2 are increasingly recognized, a major unresolved question is how these isoforms are regulated by endogenous metabolites. Small molecule intermediates such as NAD^+^, αKG, citrate, ATP, ADP, and glutamate fluctuate with nutrient status, energy demand, and redox state and have been proposed to modulate MDH activity through multiple mechanisms, including competitive inhibition, ligand binding, and effects on enzyme conformation (De Lorenzo L. et al. 2024; Raveal D.N. et al. 1962; Bernstein L.H. et al. 1978; Mueggler P.A. and Kaplan N.O. 1967; Mullinax T.R. et al. 1982; Martinez-Vaz B.M. et al. 2024; Takahashi-Iniguez T. et al. 2016; Aguero F. et al. 2004).

Despite these observations, the direct effects of these metabolites on human MDH1 and MDH2 remain poorly defined. Much of the available data derives from bacterial or plant systems, or from mammalian studies that do not distinguish between isoforms. In addition, reported regulatory outcomes vary substantially with experimental conditions, including pH, substrate and cofactor concentrations, and reaction direction. As emphasized in recent analyses (De Lorenzo L. et al. 2024; Martinez-Vaz B.M. et al. 2024), the lack of standardized experimental frameworks has made it difficult to determine how these metabolites regulate MDH activity in mammalian systems and whether the two isoforms respond differently to metabolic cues. Given MDH’s central role in cellular metabolism, the absence of a consistent framework for isoform-specific regulation by metabolic intermediates represents a gap in our understanding of mammalian metabolic control. Structural data on metabolite-bound MDH remain limited, particularly for hMDH1, and the relationship between ligand interaction, enzyme activity, and conformational dynamics in human isoforms is not well established.

To address these questions, we performed a comparative analysis of recombinant human hMDH1 and hMDH2 under matched experimental conditions. Using a combination of enzymatic assays, fluorescence-based interaction measurements, small-angle X-ray scattering (SAXS), and computational analysis, we examined how selected metabolic intermediates influence enzyme activity, ligand interaction, and structural properties. This approach allows direct evaluation of isoform-specific responses and provides a framework for understanding how local structural and dynamic features may contribute to differences in metabolite sensitivity. These findings provide a basis for interpreting how MDH isoforms may differentially respond to changes in intracellular metabolite levels and redox conditions.

## 2. Results

### 2.1 Effects of metabolic intermediates on MDH activity

To determine whether metabolic intermediates differentially regulate cytosolic and mitochondrial MDH isoforms, we measured the activity of recombinant hMDH1 and hMDH2 in the presence of ADP, ATP, citrate, α-ketoglutarate, glutamate, and NAD^+^ at concentrations of 0.1, 1.0, and 10.0 mM. Activity was normalized to the uninhibited control, which was set to 100%, and statistical comparisons were performed relative to this baseline (Figure 2). Because multiple metabolites and concentrations were tested, we emphasize concentration-dependent and isoform-specific trends rather than isolated significant values.

**Figure 2.**
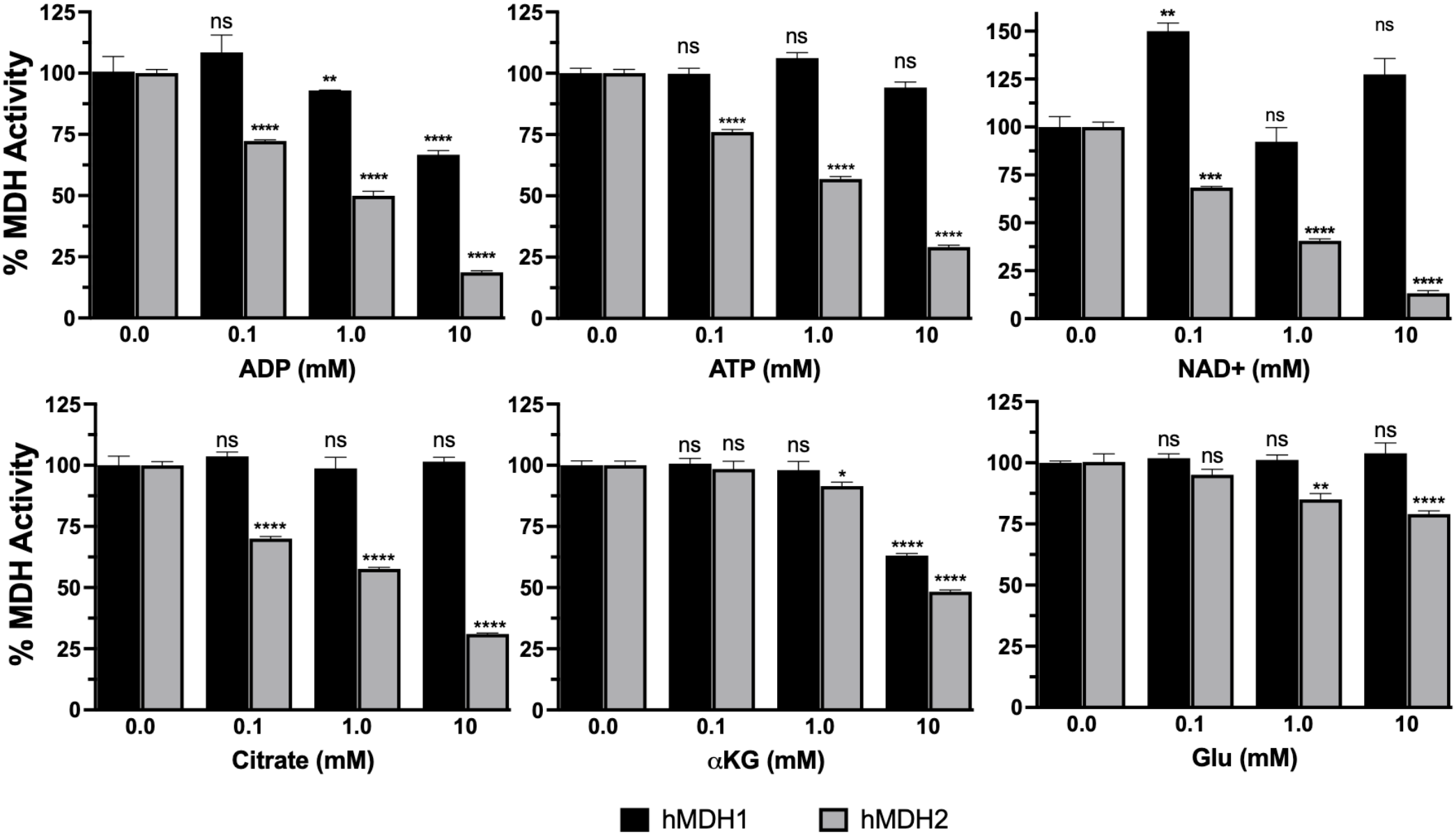
Inhibition of human MDH1 and MDH2 by central metabolic intermediates. Recombinant human cytosolic (Black bars; hMDH1) and mitochondrial (light grey bars; hMDH2) malate dehydrogenase activities were measured in the presence of ADP, ATP, citrate, αKG, Glu, or NAD^+^ at concentrations of 0.1, 1.0, and 10.0 mM. Enzyme activity was normalized to uninhibited controls (100%), and values are presented as mean percent activity ± SEM (n = 6 replicates per condition). Bars represent mean activity for hMDH1 and hMDH2 at each concentration. Statistical significance was assessed using one-sample t-tests against a normalized baseline of 100%. Annotated values above each bar indicate the *p*-value and significance level compared to control: *p* < 0.05 (*), < 0.01 (**), < 0.001 (***), and < 0.0001 (****); “ns” denotes not significant.

ADP inhibited both isoforms, but hMDH2 was more sensitive across the concentration range. hMDH1 activity was modestly reduced at 1.0 mM ADP, retaining 93.0% of control activity, and declined to 66.7% at 10.0 mM. In contrast, hMDH2 activity was significantly reduced at all ADP concentrations tested, retaining 72.3%, 49.9%, and 18.7% activity at 0.1, 1.0, and 10.0 mM, respectively.

ATP, citrate, and NAD^+^ showed stronger isoform selectivity. hMDH1 activity was not significantly affected by ATP or citrate at any concentration tested, whereas hMDH2 activity decreased progressively with increasing ATP and citrate. In the presence of ATP, hMDH2 retained 76.0%, 56.9%, and 29.1% activity at 0.1, 1.0, and 10.0 mM, respectively. Citrate produced a similar pattern, with hMDH2 retaining 70.1%, 57.7%, and 31.0% activity across the same concentration range. NAD^+^ also preferentially inhibited hMDH2, reducing activity to 68.4%, 40.6%, and 13.2% of control at 0.1, 1.0, and 10.0 mM, respectively. By comparison, hMDH1 was largely resistant to NAD^+^ inhibition and showed a small increase in activity at 0.1 mM.

Glutamate and αKG produced more limited effects but retained the same overall pattern of greater hMDH2 sensitivity. αKG affected hMDH1 only at 10.0 mM, where activity was reduced by 36.9%. In contrast, hMDH2 was inhibited at both 1.0 and 10.0 mM α-ketoglutarate, with 8.5% and 51.7% inhibition, respectively. Glutamate selectively inhibited hMDH2, reducing activity to 85.2% at 1.0 mM and 79.0% at 10.0 mM, while hMDH1 activity remained unchanged across the concentrations tested.

At 10.0 mM, several metabolite concentrations exceed typical intracellular levels and are best interpreted as testing maximal inhibitory capacity rather than direct physiological regulation. However, the consistent isoform-dependent pattern across metabolites indicates that hMDH2 is broadly sensitive to metabolic intermediates, whereas hMDH1 remains comparatively resistant under the same assay conditions.

### 2.2 Fluorescence analysis indicates selective, low-affinity interactions between MDH isoforms and metabolites

To assess whether the inhibitory effects of metabolic intermediates are associated with direct interactions, we monitored changes in intrinsic fluorescence upon ligand addition. Fluorescence emission was recorded for H108W hMDH2 and wild-type hMDH1 across increasing ligand concentrations, and apparent binding responses were estimated from concentration-dependent quenching (Table 1). This residue (H131) is surface-exposed and located away from the catalytic site, making it a suitable position for introducing a fluorescence reporter without directly modifying active-site residues. Because the two variants (wild-type hMDH1 and hMDH2 H108W) were measured using different fluorescence reporters, these values reflect fluorescence-detected interactions and should not be interpreted as direct measures of relative binding affinity between the two isoforms. Under identical assay conditions, wild-type hMDH2 and hMDH2 H108W exhibited comparable apparent turnover numbers of 13,354 ± 472 min^−1^ and 14,038 ± 1,068 min^−1^, respectively (mean ± SD, n = 6). These results indicate that introduction of the tryptophan reporter did not substantially compromise catalytic activity under the conditions examined, although effects on individual ligand-binding interactions cannot be excluded.

**Table 1.** Apparent dissociation constants (K_d_) for metabolite binding to hMDH1 and hMDH2 measured by intrinsic fluorescence quenching. Binding affinities were determined by monitoring quenching of intrinsic tryptophan fluorescence upon titration of each metabolite into solutions of recombinant hMDH1V3 and H108W hMDH2. K_d_ values (in mM) were obtained fluorescence intensity versus ligand concentration using the Benesi-Hildebrand fitting. K_d_ values reflect fluorescence-detected changes in the local environment of the reporter residues and may not capture all ligand-binding interactions. Data represent the mean ± SEM from triplicate experiments. ND = no detectable fluorescence response. Asterisks denote statistically significant differences in K_d_ between isoforms for the same metabolite, determined by Welch’s *t*-test assuming unequal variances: *p* < 0.05 (*), < 0.01 (**), < 0.001 (***); ns = not significant.

| **Ligand** | **K<sub>d</sub> binding hMDH1 (mM)** | **K<sub>d</sub> binding hMDH2 (mM)** | **Binding difference between isoform** |
|---|---|---|---|
| α-ketoglutarate | 18.9 ± 1.45 | 10.4 ± 1.50 | <sup>\*\*</sup> *p=0.0018* |
| Citrate | ND | ND |  |
| Glutamate | ND | ND |  |
| ATP | 16.1 ± 1.80 | 9.63 ± 0.68 | \* *p=0.015* |
| ADP | 4.61 ± 0.63 | 3.77 ± 0.27 | ns *p=0.134* |
| NAD<sup>+</sup> | 1.95 ± 0.01 | 1.02 ± 0.04 | \*\*\* *p=0.001* |

NAD^+^ produced the largest fluorescence response in both isoforms, consistent with measurable low-millimolar interactions. The apparent affinity was higher for hMDH2 (K_d_ ≈ 1.0 mM) than for hMDH1 (K_d_ ≈ 2.0 mM), indicating modest but reproducible isoform selectivity. ADP also produced clear fluorescence changes in both isoforms, with similar apparent affinities in the mid-millimolar range (approximately 3–5 mM).

ATP and α-ketoglutarate generated weaker responses overall but retained a consistent pattern of stronger interaction with hMDH2. Apparent affinities for these ligands were in the high-millimolar range, with ATP and α-ketoglutarate binding hMDH2 at approximately 9–10 mM and hMDH1 at approximately 15–20 mM. These values indicate relatively weak interactions that are only partially aligned with the concentrations required for enzymatic inhibition.

Citrate and glutamate did not produce detectable fluorescence changes under the conditions tested. Because this assay reports changes in the local environment of the reporter residue, the absence of signal does not exclude interaction but suggests that any interaction is either weak, transient, or not coupled to the fluorescence probe. The engineered tryptophan in hMDH2 is positioned near the nucleotide-binding region and therefore preferentially reports interactions in that local environment.

Overall, these results indicate that only a subset of metabolites produce detectable interactions in this assay and that these interactions are generally low affinity. The consistently stronger fluorescence responses observed for hMDH2 suggest greater responsiveness to metabolite binding. However, the relatively weak apparent affinities and the use of different fluorescence reporters limit direct comparisons between the isoforms and suggest that fluorescence-detected interactions represent only a subset of the full interaction landscape.

### 2.3 SAXS analysis reveals stable dimeric structure and isoform-specific flexibility without ligand-induced global rearrangement

As of this writing, there are 100+ crystal structures of malate dehydrogenases in the PDB, all showing a similar dimeric or tetrameric structure. Ligand binding to the protein causes minute changes in the conformation of the protein, primarily in the active site loop, which binds to the dicarboxylic acid substrate and leads to hydride transfer [5]. However, it remains unclear from crystallographic data how metabolites such as citrate, αKG, and nucleotide compounds, known to modulate MDH activity, affect the overall conformation of MDH or influence function through structural mechanisms. To explore these questions, we used SAXS to investigate whether these small molecules induce conformational changes in solution that are not captured in static crystal structures.

Given the previous X-ray crystallography data showing minute changes in the MDH active site structure induced by ligand binding, we chose to use solution small-angle X-ray scattering (SAXS) to observe if these molecules regulated activity by altering the overall structure of the MDH dimer. *E. coli* MDH is known to dissociate reversibly under certain conditions to inactive monomers [16] and has subsequently been shown for porcine mitochondrial MDH upon dilution and at low pH, whereas MDH1 does not appear to undergo the same dissociation [17,18]. Given the isoform-selective nature of this phenomenon, and the difference in isoform responsiveness to metabolite binding and inhibition, assessing whether ligand binding or oligomeric state differs between human isoforms of MDH in solution is worth investigating and well addressed by SAXS analysis. Data on the wild-type MDH2 dimer showed a molecular weight of a 62.4 kDa and an R_g_ of 28.6 ± 0.1 Å (Table 2). When we fit the SAXS data to the hMDH2 crystal structure (2DFD) using FOXS, we observed a reasonable match between a model of the MDH2 protein and the SAXS data (X^2^ = 1.1). This is the first solution characterization of hMDH2 (Figure 3) and the data indicate that the crystallographic models match the solution conformation of MDH within the resolution of SAXS.

**Table 2.**
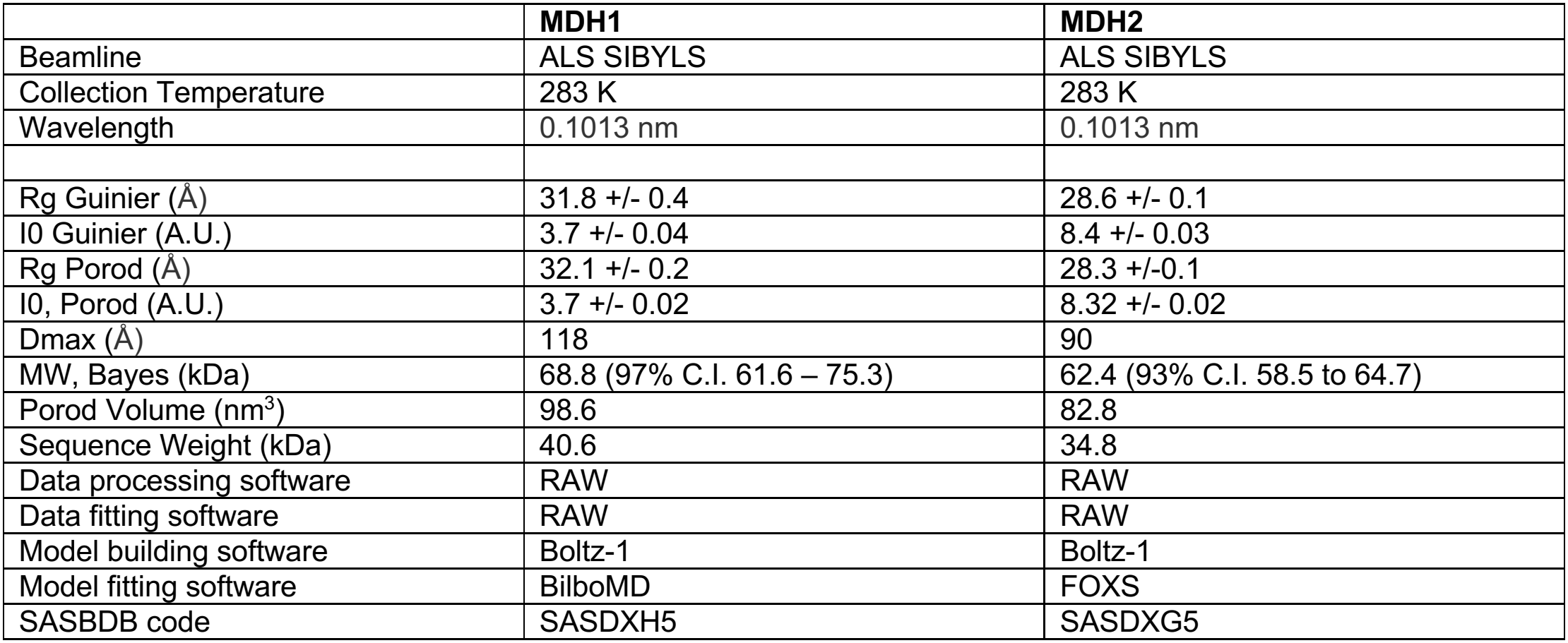
SAXS parameters.

**Figure 3.**
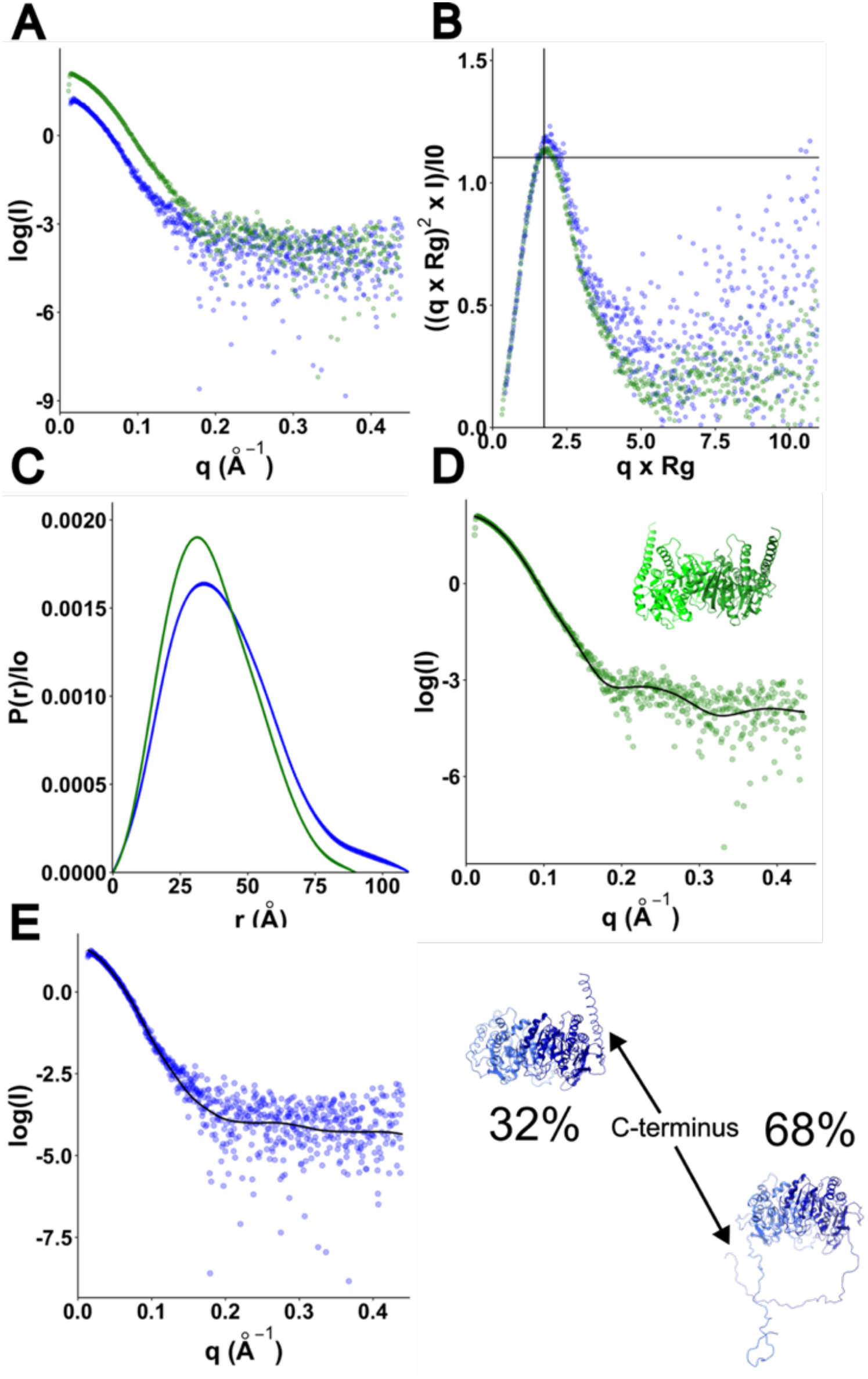
SAXS data and fitting. (A) Log of intensity vs. q (A^-1^) for MDH1 (blue) and MDH2 (green) (B) normalized Kratky replots of intensity (C) paired distance distribution function (PDDF) plots with error bars. For the PDDF fitting, the X^2^ values were 0.997 and 1.092 and total estimates of 0.865 (Good) and 0.952 (Excellent) for MDH1 and MDH2, respectively. (D) FOXS of the Boltz-1 model of MDH2 to the SAXS data. The X^2^ = 1.1 (E) BilboMD fit of MDH1 model from Boltz-1 to the SAXS data. Using a two-state model improved the X^2^ from 1.13 to 1.07. The single state model with slightly poorer fitting is shown in the bottom right of the figure.

For comparison, we collected SAXS data on hMDH1, which also lacks a complete solution characterization. MDH1 had an R_g_ of 31.8 ± 0.4 Å and a molecular weight of 80.1 kDa. Fitting the crystal structure of hMDH1 5NUF to the data produced an X^2^ of 1.98, with a reasonable match at low q values but a poorer fit in the Porod and wide q region. To improve the modeling, we then modeled the structure of the expressed sequence of an hMDH1 dimer in Boltz-1 [19], resulting in a model with a confidence score of 0.9, which suggests a highly likely model. We then fitted the resulting dimer to the SAXS data, resulting in an X^2^ of 1.34, which is better than the crystal structure but still suffered from poor fitting at wide q values. We refined the model further using BilboMD, which performs limited MD on structures to fit the SAXS data, finding that unfolding the C-terminal helix improved the X^2^ to 1.17. Using a two-state model with a mix of folded and unfolded C-termini decreased the X^2^ to 1.07. Adding additional conformations did not improve the fit. Thus, we find it likely that hMDH1 has a flexible C-terminal region that adopts a disordered structure at least part of the time. This C-terminal flexibility is missing from the existing crystal structures of hMDH1

A comparison of the SAXS data on the two human MDH proteins indicates that both proteins are well-folded dimers with similar shapes. However, MDH1 is slightly larger based on the larger R_g_ and D_max_ values (Table 2). Comparison of the P(r) plots suggests that the core region of the dimer is similar in size and dimensions, as indicated by the similar distance value of the plot peak, which supports the similarity of the crystal structures (5NUF and 2DFD and Figure 3). However, the longer dimensions observed with lower probability on the right side of the plot are due to the C-terminal unfolding in hMDH1.

To test whether the binding of the ligands resulted in global changes to the structure of hMDH1 or hMDH2, we collected SAXS data on both proteins in the presence of ligands. These data suggest that neither isoform undergoes significant structural rearrangement or oligomer dissociation upon ligand binding, and that metabolite effects on activity likely arise from local or dynamic perturbations rather than global conformational changes.

The lack of an observable conformational change in either hMDH1 or hMDH2 next led us to explore potential differences in protein dynamics. We observed that the active-site loop and a dynamic loop adjacent to it (“thumb”) have different sequences and conformations in crystal structures [5]. Alignment of the crystal structures of MDH1 and MDH2 enzymes in the RCSB PDB shows that the diversity of active-site loop conformations in MDH1 enzymes is greater than that in MDH2 enzymes, which are all in the closed state (data not shown). The sequence of hMDH1’s active-site loop carries a net charge of +2 (six charged residues: four positive and two negative) compared to +1 in hMDH2 (five charged residues: three positive and two negative); hMDH2 also carries an insertion in the thumb loop, extending it by two amino acids relative to hMDH1 (Figure 4A). Highly charged protein sequences are generally dynamic, and adding two negatively charged side chains to the loop could enhance dynamics and destabilize anion binding at the active site. To identify the potential effects of these substitutions, we performed explicit-solvent molecular dynamics simulations on the modeled structures, which we supported with normal mode analysis, both of which can predict the structural dynamics of each protein [20,21]. Focusing on both proteins’ active sites shows that the active site of hMDH1 is more dynamic than hMDH2, while the thumb loop is similarly dynamic. Given that the active site loop seems to close to hold the dicarboxylic substrate in place during catalysis, the increased dynamics of the active loop and thumb loop in hMDH1 may lead to higher specificity, requiring more specific contacts before the loop closes in a stable position. Substrates or potential inhibitors that don’t fit as well in the active site are less likely to be “captured” in the active site, leading to catalysis. We can further infer that the lower catalytic rates for MDH1 may also be a consequence of the higher specificity and dynamics.

**Figure 4.**
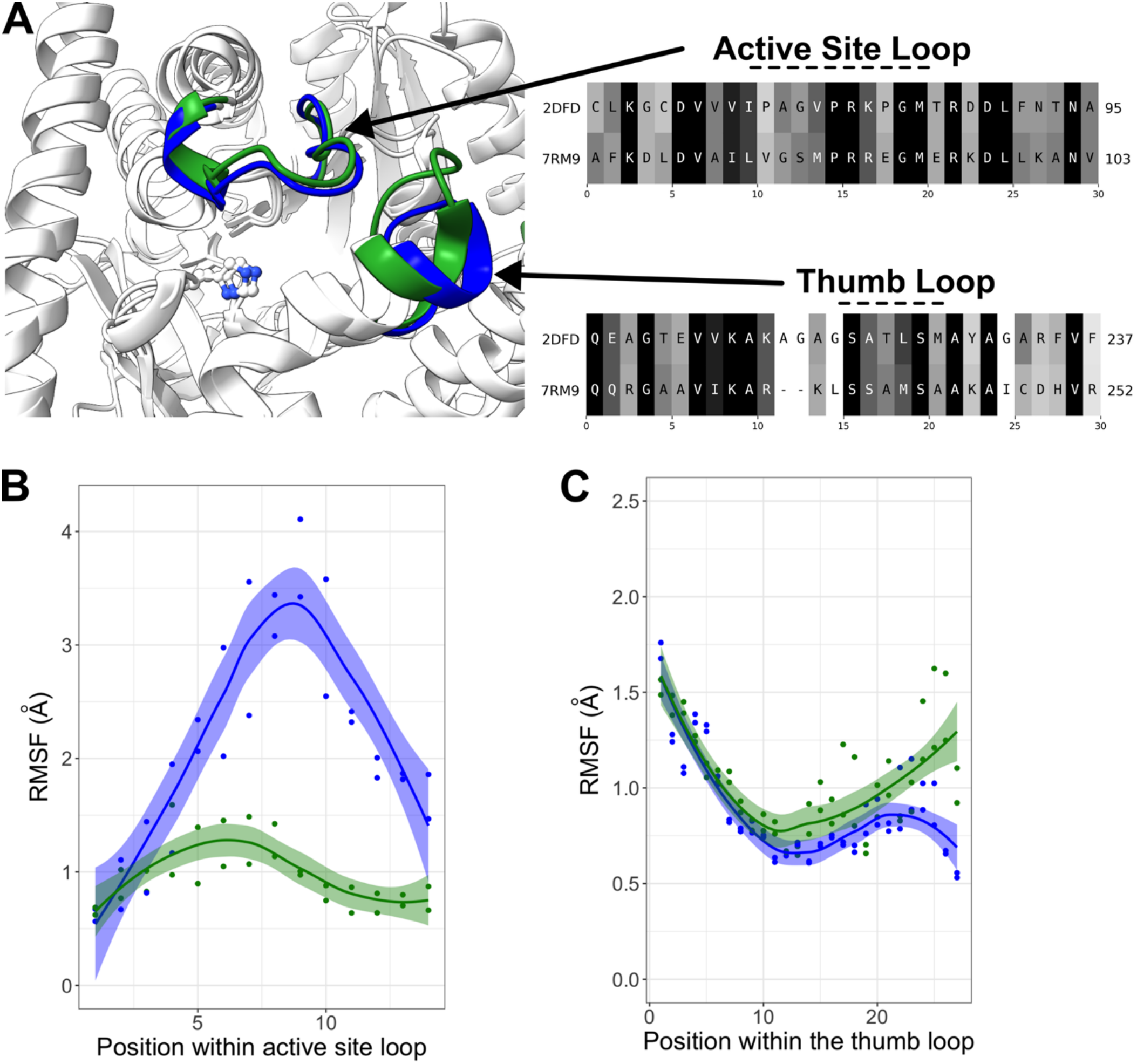
Dynamics of the MDH active site loops. (A) Structural alignment of the active site loop and thumb for MDH1 (blue, PDB ID: 7RM9) and MDH2 (green, PDB ID: 2DFD), RMSD of 1.2 Å. The sequence alignments were extracted from the structural alignment using the RCSB Structural Alignment tool. (B) Root Mean Square Fluctuation (RMSF) of MDH1 (blue) and MDH2 (green) as calculated from the MD trajectory for the active site loop (B) and the thumb loop (C).

An additional aspect of substrate binding is the active site environment. We assessed the active environment using PropKa3 to calculate the pKa values of the catalytic histidine [22]. For hMDH1, the range of catalytic histidine pKa values during the trajectory was 4.7 to 8.3 with a mean value of 7.6 +/- 0.3. For hMDH2, the pKa range was 2.2 to 7.7 with a mean value of 5 +/- 1.1. We next correlated the active site Histidine pKa value with the distance from the loop and noticed that closure of the hMDH2 loop results in a more drastic decrease in the pKa value. The data for hMDH1 show a similar trend; however, the decrease in pKa is smaller (1 pH unit versus 2-4 units). These data suggest that the chemical environment of the hMDH2 active site is more diverse than the hMDH1 active site. We then compared the pKa values for the human, ligand bound crystal structures of MDH1 and MDH2 in the PDB (Figure 5C). For structures of MDH2, the the unbound or cofactor-only boudn structures match our predictions from the MD of the apo MDH2, with a predicted His pKa value of ∼5. Once a carboxylate substrate or inhibitor is bound, the His pKa increases to ∼7.5 to 8. For MDH1, which has fewer structures in the database, we observe the apo form has predicted His pKa values of 7.4 and 5.4, while the subunits in the malonate bound form has predicted pKa values of ∼7.5. The wide difference in values for the apo form, we believe is due to misalignment of the His-Asp pair in the B subunit (Figure 5D). Prediction of pKa values for the active site Asp showed smaller to no increases in pKa values depending on the ligand bound state, suggesting this position is less sensitive to the active site state. Collectively, these data support the inhibition studies showing that hMDH2 is broadly inhibited by ligand molecules while hMDH1 shows minimal inhibition. We link these differences in molecule binding to the slower active site loop dynamics of hMDH2, slowing inhibitor dissociation from the active site, and the diverse hMDH2 active site environment, allowing more molecules to “fit”.

**Figure 5.**
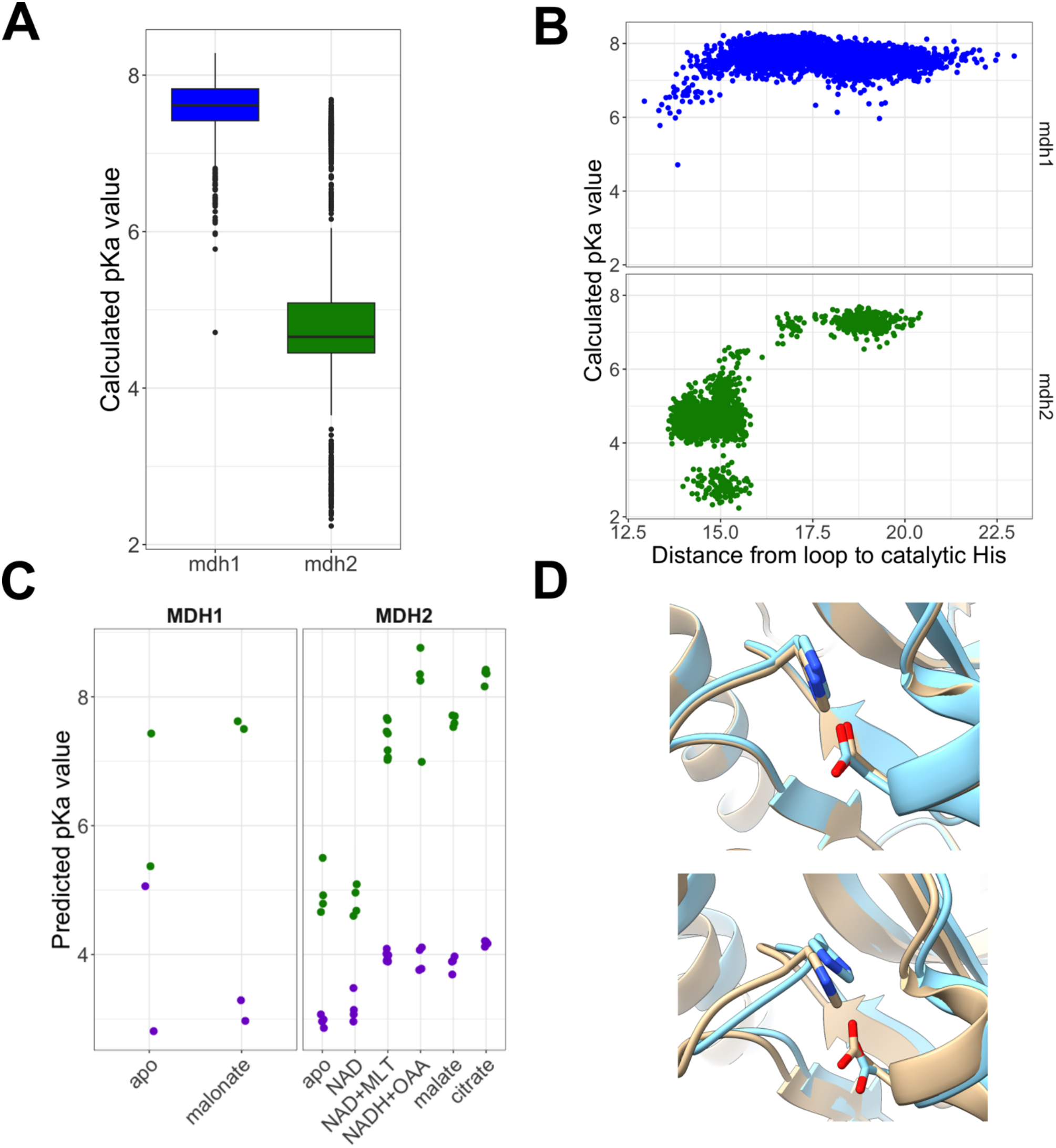
Chemical environment of MDH active site. (A) Predicted pKa values of the active site Histidine from MD calculated from the MD trajectory. (B) Effect of active site loop position on pKa of the active site. (C) PROPKA predicted pKa values from crystal structures with ligands bound. Crystal structures for MDH1 with 7RRL and 7RM9 and for MDH2, 2DFD, 4WLE, 4WLF, 4WLN, 4WLO, 4WLU, and 4WLV. The pKa values of the active site Histidine are shown in green, while for the values for the active site aspartate are shown in purple. (D) Orientation of the active site His-Asp pair for MDH1 as shown in structure 7RRL (tan) and 7RM9 (cyan) for the A (top panel) and B (bottom panel) subunits.

## 3. Discussion

Although hMDH1 and hMDH2 catalyze the same NAD^+^-dependent redox reaction, their compartmentalization and structural differences suggests that they may be subject to distinct regulatory control. Here, we evaluated the effects of multiple metabolic intermediates on enzyme activity, fluorescence-detected ligand interactions, and solution structure using a unified experimental framework. This approach allows direct comparison of isoform behavior under matched conditions and reduces variability associated with prior studies using different species or assay systems. Prior studies of MDH regulation have frequently utilized non-human or microbial enzymes or have applied varied experimental conditions that may contribute to inconsistent findings, particularly regarding the effects of citrate, ATP, and NAD^+^ (Martinez-Vas. et al. 2024). In this study, we applied a unified experimental approach to evaluate metabolite-specific regulation of the human cytosolic and mitochondrial MDH isoforms

The enzymatic data show clear isoform-specific differences in sensitivity to metabolite perturbation. hMDH2 was inhibited by multiple metabolites, including ADP, ATP, citrate, α-ketoglutarate, and NAD^+^, whereas hMDH1 maintained activity across a wider range of conditions. Overall, this pattern indicates that the mitochondrial isoform is more responsive to metabolite perturbation than the cytosolic isoform.

Previous studies have reported variable effects of metabolic intermediates on MDH activity, often dependent on isoform, pH, and assay conditions. Citrate, for example, has been described as both an activator and inhibitor across different systems (Genda T. et al. 2003; Mullinax T.R. 1982; Wise D.J. et al. 1997; Bell J.K. et al. 2001). Under the matched conditions used here, citrate inhibited hMDH2 in a concentration-dependent manner, whereas hMDH1 activity was not significantly effected. Glutamate has been reported to activate mitochondrial MDH at alkaline pH (Fahien L.A. et al. 1998); in contrast, at pH 7.4 we observed only modest inhibition even at the highest concentrations tested. The effects of α-ketoglutarate have also been shown to depend on cofactor availability and enzyme context (Fahien L.A. et al. 1998). In our assays, α-ketoglutarate alone inhibited MDH1 only at higher concentrations where MDH2 showed inhibition at both 1 and 10 mM. Adenine nucleotides reported to inhibit MDH in both plant and animal systems (Raveal D.N. et al. 1962; Mueggler P.A. and Kaplan N.O. 1967; Takahashi-Iniguez et al. 2016). Consistent with these reports, we show that ADP inhibited both isoforms, whereas ATP inhibited hMDH2 without significantly affecting hMDH1 activity under the conditions tested. The pronounced inhibition of hMDH2 by NAD^+^ may reflect isoform-specific differences in product inhibition or in the relative occupancy of nucleotide-bound enzyme states. The modest increase in hMDH1 activity observed at 0.1 mM NAD^+^ may reflect a different response to nucleotide occupancy, although the present experiments do not establish the mechanism underlying this effect. Further kinetic analysis varying both NADH and NAD^+^ concentrations would be required to distinguish these possibilities.

The relationship between inhibition and fluorescence responses places important constraints on interpretation. Metabolites such as NAD^+^ and ADP produced both measurable fluorescence changes and inhibition, whereas citrate and glutamate inhibited hMDH2 activity without producing detectable fluorescence responses under the conditions tested. This indicates that fluorescence-detected perturbation and functional inhibition are not equivalent readouts. Because the fluorescence assay reports changes in the local environment near the probe, it likely captures only a subset of interactions. Taken together, these data support low-affinity, fluorescence-detectable interactions for some metabolites, while indicating that additional inhibitory effects may arise from weak, transient, or spatially remote interactions not reported by the fluorescence probe. These data do not distinguish between active-site binding, alternative binding sites, or indirect effects on the assay environment.

The SAXS data further constrain the structural basis of inhibition. Both isoforms exist as stable dimers in solution, and no evidence of large-scale conformational rearrangement or oligomer dissociation was observed upon ligand addition. These findings indicate that metabolite-dependent inhibition is not associated with global structural transitions detectable at the level of overall molecular shape. Instead, the inhibitory effects appear to occur within a structurally stable dimeric framework.

Structural modeling and computational analyses provide additional context for the observed isoform differences. SAXS-guided modeling is consistent with hMDH1 sampling a broader conformational ensemble than hMDH2, including partial flexibility in the C-terminal region. Normal mode analysis further suggests increased mobility in the active site loop and adjacent regions of hMDH1 relative to hMDH2. These observations are consistent with differences in sequence and loop architecture between the isoforms, including modest differences in charge distribution within the active site loop of hMDH1 and a structural extension in the hMDH2 thumb loop.

Based on these combined observations, we propose a working model in which isoform-specific differences in local structure and dynamics influence metabolite sensitivity. In this model, the more constrained and comparatively rigid architecture of hMDH2 facilitates interactions with negatively charged metabolites, resulting in broader sensitivity to inhibition. In contrast, the increased flexibility and altered charge distribution in hMDH1 may reduce the stability or residence time of weakly interacting ligands, thereby limiting inhibitory effects. This interpretation is consistent with both the enzymatic and fluorescence data but does not define a specific binding mechanism.

Estimated intracellular concentrations of these metabolites vary across compartments and generally fall within micromolar to low-millimolar ranges (Table 3). Inhibition of hMDH2 was observed across part of this concentration range for several metabolites, whereas hMDH1 remained comparatively resistant. The affinities inferred from fluorescence measurements fall within the low-to mid-millimolar range, indicating relatively weak interactions that may depend on local metabolite concentrations, compartment-specific accumulation, or transient microenvironments rather than bulk cellular levels. While the present study does not directly address these conditions, the observed differences between isoforms are consistent with their distinct subcellular environments and functional roles. These findings suggest that hMDH2 may be more responsive to changes in local metabolite concentrations, whereas hMDH1 remains comparatively resistant under the conditions examined.

**Table 3.** Estimated physiological concentration ranges of MDH-regulating metabolites in mammalian cells. Estimated physiological concentration ranges of metabolites that modulate MDH1 and MDH2 activity in mammalian cells. Values are drawn from NMR, LC-MS/MS, isotope tracing, and biosensor studies. Concentrations vary based on tissue type, culture conditions, metabolic state, and analytical method. These include mass spectrometry, enzyme-coupled assays, and real-time biosensor quantification.

| **Metabolite** | **Cytosolic Concentration (µM)** | **Mitochondrial Concentration (µM)** | **Whole-Cell Concentration (µM)** |  |
|---|---|---|---|---|
| **ADP** | 130–600 | 60–2100 | 100–200 | Chen et al. 2016; Stern et al 2023; Traut et al. 1994; Soboll et al. 1980 |
| **ATP** | 1200–5000 | 20–2100 | 1200–5000 | Chen et al. 2026; Stern et al. 2023; Traut et al. 1994; Soboll et al. 1980 |
| **NAD<sup>+</sup>** | 470–1000 | 770–3900 | 750–1000 | Chen et al. 2026; Stern et al. 2023; Traut et al. 1994; Braidy et al. 2011 |
| **NADH** | 60–100 | 7–2100 | 80–100 | Chen et al. 2026; Stern et al 2023; Hung et al 2009 |
| **α-Ketoglutarate** | 300–1300 | 15–200 | 200–300 | Chen et. al. 2026,;Stern et. al 2023 |
| **Citrate** | 150–200 | 440–3700 | 150–200 | Stern et. al 2023; Zhao et. al. 2023 |
| **Glutamate** | 1000–5000 | 2000–10000 | 2000–5000 | Traut 1994; McKenna 2007; Braha et. al. 1982 |

## 4 Conclusion

Taken together, these findings demonstrate that human MDH1 and MDH2, despite their conserved catalytic mechanisms and overall structural similarity, differ substantially in their sensitivity to metabolic intermediates. These differences suggest that the regulation of MDH activity depends not only on enzyme localization and catalytic properties, but also on how each isoform responds to its metabolic environment. The fluorescence and structural analyses provide possible explanations for these differences, although the specific mechanisms of inhibition remain to be established. Future studies examining metabolite interactions, local structural dynamics, and the effects of physiological metabolic conditions will help determine how these differences contribute to the regulation of cytosolic and mitochondrial metabolism. Understanding these isoform-specific responses may provide a more complete picture of how MDH activity is coordinated with cellular energy and redox balance.

## 5 MATERIALS AND METHODS

### 5.1 Recombinant MDH cloning and purification

hMDH1 splice variant 3 (UniProt #P40925-3) was isolated from a human brain cDNA library and cloned into a pET28 vector. hMDH2 (UniProt #P40926-1) was synthesized with codon optimization for expression in *Escherichia coli*. For both constructs, a TEV protease recognition site and 6×His tag were included at the C-terminus. For fluorescence measurements, hMDH2 was modified to include a tryptophan residue (H108W). Site-directed mutagenesis was performed using QuikChange (Agilent Technologies) with the following primers: 5′-CGCTGCTTGCGCTCAGTGGTGTCCGGAAGCTATG-3′ and 5′-CATAGCTTCCGGACACCACTGAGCGCAAGCAGCG-3′. Recombinant proteins were expressed in BL21(DE3) cells grown in either 2×YT or LB media. Cultures were induced with 1 mM IPTG and incubated overnight at 20°C prior to harvest. Cells were lysed and clarified lysates were subjected to Ni-agarose chromatography. Proteins were eluted using buffer containing 10 mM Tris-Cl, 50 mM NaCl, 300 mM imidazole, and 0.1 mM EDTA, pH 7.4. Eluted fractions were pooled and dialyzed against 10 mM Tris-Cl, 50 mM NaCl, and 0.1 mM EDTA, pH 7.4. Protein purity was assessed by SDS-PAGE and was greater than 90%. Protein concentrations were determined using the Bradford assay and maintained between 0.5 and 1.5 mg/mL for subsequent experiments.

### 5.2 Enzymatic analysis

Recombinant hMDH1 and hMDH2 enzyme activity was measured spectrophotometrically at 340 nm by monitoring NADH oxidation under steady-state initial velocity conditions in the direction of oxaloacetate to malate conversion. Assays were performed at 20°C in buffer containing 100 mM potassium phosphate, pH 7.4 with 0.2 mM OAA (made fresh within 3 h of the assay) and 0.1 mM NADH in a 96-well plate format initiating the assay with injection of OAA. For each condition, activity was measured in the presence of one of six small molecule metabolites: ADP, ATP, citrate, αKG, Glu, or NAD^+^, each tested at 0.1, 1.0, and 10.0 mM final concentrations. Specific activities were calculated from the initial rates of NADH oxidation and the amount of enzyme present in each reaction. Control reactions with no added inhibitor were also performed. All measurements were performed in 4-10 technical replicates per condition and isoform. Enzyme activity was normalized to the mean activity of the no-inhibitor control for each isoform, which was defined as 100%. Data are expressed as percent activity relative to this baseline. Tests were performed separately for hMDH1 and hMDH2 for each metabolite. Statistical significance of metabolite-induced effects was evaluated using a one-sample t-test comparing each condition to the normalized control value (100%). Tests were performed independently for each metabolite and isoform. p-values were reported alongside effect sizes and interpreted using standard thresholds (p < 0.05, < 0.01, < 0.001, < 0.0001). All analyses were performed using GraphPad Prism.

### 5.3 Comparison of wild-type MDH2 and MDH2 H108W

Apparent turnover numbers for hMDH2 H108W and wild-type hMDH2 were determined by monitoring NADH absorbance at 340 nm. Assays were performed in 100 mM phosphate buffer containing NADH and oxaloacetate prepared fresh in deionized water and maintained on ice prior to use. Reaction mixtures contained final concentrations of 100 mM phosphate buffer, 0.1 mM NADH, and 0.5 mM oxaloacetate. Enzyme was added at a concentration sufficient to produce linear, first-order changes in absorbance over a 20–30 s interval. Reactions were initiated by addition of enzyme and monitored immediately using a Cary 60 UV–vis spectrophotometer. Initial rates were determined from the linear portion of the absorbance trace, and values from six independent measurements were averaged. Apparent turnover numbers were calculated by converting ΔA340/min to concentration using the molar extinction coefficient for NADH (ε340 = 6220 M^−1^ cm^−1^) and dividing by enzyme concentration. Errors were calculated from the standard deviation of replicate measurements.

### 5.4 K_d_ determination of hMDH2 H108W and hMDH1

Changes in intrinsic fluorescence emission upon ligand addition were used to estimate apparent dissociation constants (K_d_,app) using double-reciprocal plots derived from the Benesi–Hildebrand relationship (Benesi and Hildebrand, 1949). Fluorescence measurements were performed in 96-well plates containing fixed concentrations of recombinant protein and varying concentrations of ligand in 100 mM potassium phosphate buffer, pH 7.4. Ligand concentrations ranged from 2–14 mM for ATP and NAD^+^, 2–7 mM for ADP, and 2–12 mM for αKG with hMDH2 H108W or 6–14 mM with hMDH1. No measurable fluorescence quenching was detected with sodium citrate at concentrations up to 1.33 M, glutamate up to 850 mM, or glutamine up to 150 mM. Fluorescence was measured using a SpectraMax M2e multimode microplate reader (Molecular Devices). Samples were excited at 290 nm, and emission was recorded from 310 to 400 nm in 10 nm increments. The photomultiplier tube (PMT) gain was set to high for hMDH1 and 600 V for hMDH2 H108W. Fluorescence intensity at 340 nm was used for quantitative analysis. Each ligand titration was performed with three replicate measurements at each concentration, and the complete experiment was repeated independently three times. Fluorescence intensities from a representative experiment are presented as mean ± SD in Supplementary Figure S1, together with the corresponding Benesi–Hildebrand analyses. Apparent dissociation constants reported in Table 1 were obtained from the original experimental analyses. Assuming a 1:1 binding stoichiometry, the Benesi–Hildebrand relationship was expressed as: 1/ΔF = 1/((F_∞_-F_o_).K_d_[L]) + 1/(F_∞_-F_o_), where F_o_ is the fluorescence intensity in the absence of ligand (L), F_∞_ is the fluorescence intensity with saturating concentration of ligand, F is the fluorescence intensity at some particular [ligand], DF is (F-F_o_), and K_d_ is the dissociation constant. When 1/DF is plotted against 1/[ligand], a linear relationship exists. The intercept on the x-axis is –1/K_d_. The data are reported with error determined at the 95% confidence level

### 5.5 SAXS analysis

hMDH1 and hMDH2 were purified with a HisTrap HP 1 mL Ni column using an AKTA Start FPLC. Cells were lysed in 50 mM Tris-HCl, pH 8.0, 1 mM imidazole, 100 mM NaCl, and 0.1 mM EDTA. The column was washed with 20 mL of 50 mM Tris HCl, pH 8.0, 300 mM NaCl, 10 mM imidazole, 0.1 mM EDTA and eluted with a linear gradient of the wash buffer and the elution buffer 50 mM Tris-HCl, pH 8.0, 300 mM Imidazole, 50 mM NaCl, and 0.1 mM EDTA. Fractions with MDH activity were pooled and run through a Sephacryl S100 size exclusion column to remove aggregated and degraded polypeptides. The size-exclusion buffer consisted of 50 mM Tris-Cl,pH 8.0, 100 mM NaCl, and 0.1 mM EDTA. Pure samples were identified via SDS-PAGE. Samples for SAXS were dialyzed into 100 mM potassium phosphate, pH 7.4. SAXS samples were loaded onto a SAXS plate at concentrations ranging from 0.5 to 8 mg/mL. The samples and size-exclusion buffer used as a protein-free sample were frozen at -80ºC before shipment. SAXS data were collected on SIBYLS beamline 12.3.1 at the Advanced Light Source at Lawrence Berkeley National Laboratory (Classen S et al. 2013 and Rosenberg DL et al. 2022) Before sample collection, the plate was spun at 3700 rev min^-1^ for 10 minutes. Samples were held at 10 ºC during collection. The exposure was 15 seconds, with frames collected every 0.3 seconds for 40 frames per sample. The incident light wavelength was 1.03 Å at a sample-to-detector distance of 2 m. This setup results in scattering vectors, *q*, ranging from 0.013 to 0.5 Å^™1^, where the scattering vector is defined as *q* = 4πsin*θ*/*λ*, and 2*θ* is the measured scattering angle. Radially averaged data were processed using RAW 2.2 (Hopkins JB et al. 2024) to determine the protein dimensions. Structural models and crystal structures were fitted to SAXS data using FOXS (Schneidman-Duhovny D et al. 2016). The model of hMDH1 was refined and fitted to the SAXS data using BilboMD (Pelikan M et al. 2009). SAXS data and the fitted models for MDH1 and MDH2 are available on the SASBDB under codes SASDXH5 and SASDXG5, respectively (Kikhney AG et al. 2019).

### 5.6 Molecular Dynamic Analysis of Small Molecule Binding

Models of the expressed sequences of hMDH1 or hMDH2 were modeled using Boltz-1 (Boltz-1; Colabfold; Wohlwend J et al 2024 and Kim G et al. 2024) running in Google Colab. To model ligand binding, we further provided ligands as SMILES strings. Ternary complexes were modeled by providing two ligands to the algorithm with NADH in the second position. Thus, the protein and ligands were modeled simultaneously. Positions of the ligands were visualized in ChimeraX (Pettersen EF et al. 2021). For normal mode analysis of hMDH1 and hMDH2, we used the Bio3D package (ver 2.4-5) in R (Grant BJ et al. 2006). The Boltz models of both proteins were provided as PDB files, the PDB files aligned, and the normal modes simulated in Bio3D. We then compared the resulting root mean squared fluctuations (RMSF) from simulated normal modes. Active site pKa values were calculated using PROPKA running in Google Colab (Olsson MHM et al. 2011).

## DATA AVAILIBILITY STATEMENT

SAXS data and fitted structural models for hMDH1 and hMDH2 have been deposited in the Small Angle Scattering Biological Data Bank (SASBDB) under accession codes **SASDXH5** and **SASDXG5**, respectively. All processed enzyme activity data and calculated inhibition values are provided within the article and its supporting information. Data are available upon request.

## Supplemental Data

**Supplementary Figure S1.**
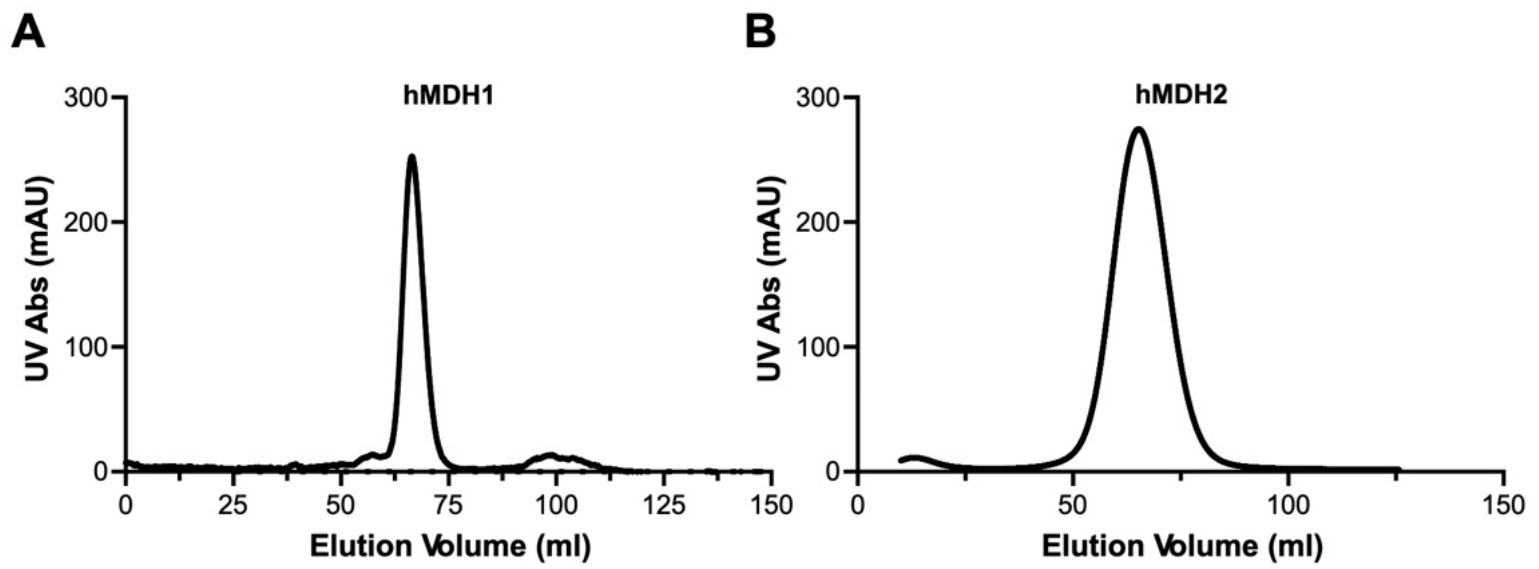
Size-exclusion chromatography of recombinant human MDH isoforms. Following imidazole elution from affinity chromatography, recombinant hMDH1 (A) and hMDH2 (B) were concentrated using YM-10 Centricon centrifugal concentrators (Fisher Scientific) and subjected to size-exclusion chromatography on a Sephacryl S-200 column equilibrated with 10 mM potassium phosphate (pH 7.4), 50 mM NaCl, and 0.1 mM EDTA. Fractions corresponding to the major elution peaks were pooled and assessed for protein purity by SDS-PAGE.

**Supplementary Figure S2.**
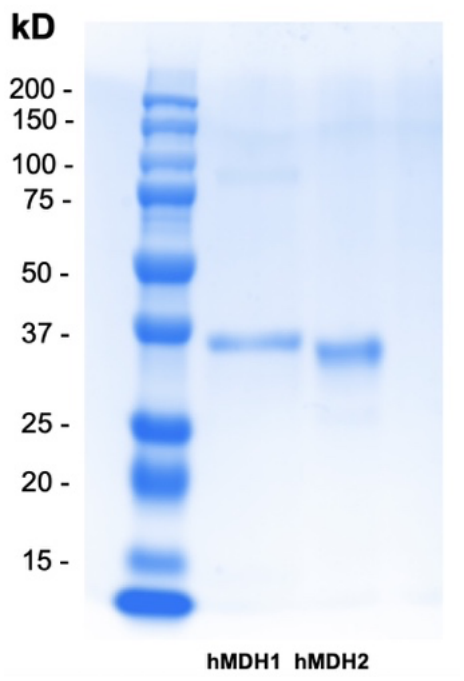
SDS-PAGE analysis of purified recombinant human MDH isoforms. Pooled fractions from Sephacryl S-200 size-exclusion chromatography were analyzed by SDS-PAGE using a 12% polyacrylamide gel (Bio-Rad). Protein bands were visualized by Coomassie staining. Lane 1, broad-range molecular weight standards (Bio-Rad); lane 2, recombinant hMDH1 (3 µg); lane 3, recombinant hMDH2 (3 µg).

**Supplementary Figure S3.**
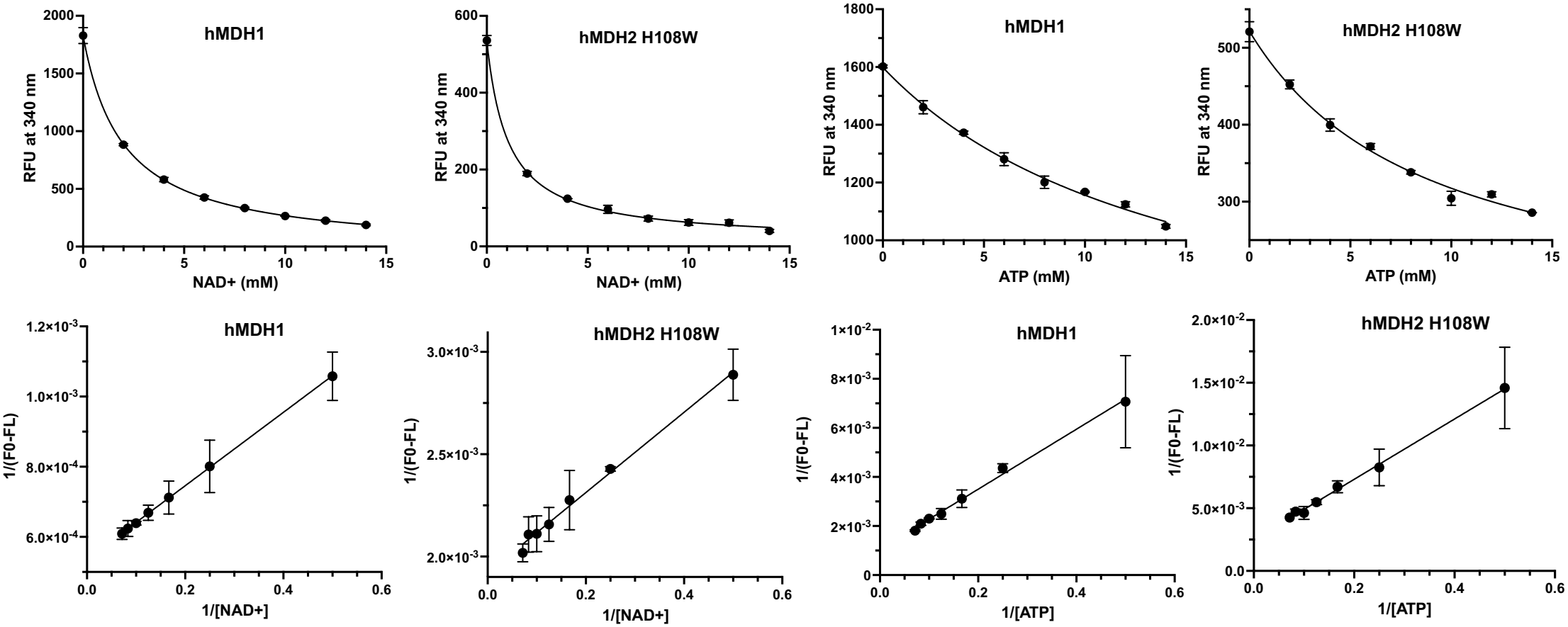
Fluorescence titration and Benesi–Hildebrand analysis of NAD^+^ and ATP interactions with human MDH isoforms. Concentration-dependent changes in intrinsic fluorescence were measured for recombinant hMDH1 and H108W hMDH2 following addition of NAD^+^ or ATP. Top panels show fluorescence intensity at 340 nm as a function of ligand concentration for hMDH1 with NAD^+^, H108W hMDH2 with NAD^+^, hMDH1 with ATP, and H108W hMDH2 with ATP. Data are presented as mean ± SD of three replicate measurements from one representative experiment. Each experiment was performed independently three times. Panels in the bottom row show the corresponding Benesi–Hildebrand double-reciprocal plots of 1/|ΔF| versus 1/[ligand], with linear regression used to estimate apparent dissociation constants (K_d_,app), assuming a 1:1 binding relationship. Fluorescence measurements were performed in 100 mM potassium phosphate buffer, pH 7.4, using excitation at 290 nm. The representative titrations illustrate the concentration-dependent fluorescence responses and analytical approach used to determine the apparent affinities reported in Table 1.

During the preparation of this manuscript, the authors used ChatGPT (OpenAI) to assist with manuscript organization, and clarity. The authors reviewed and edited the output and take full responsibility for the content of the published article.

## REFERENCES

1. Aguero F, Noe G, Hellman U, Repetto Y, Zaha A, Cazzulo JJ. Purification, cloning, and expression of the mitochondrial malate dehydrogenase from protoscolices of Echinococcus granulosus. Mol Biochem Parasitol. 2004;137:207–214.

2. Braha S, Clarke DD. Cerebral glutamine/glutamate interrelationships and metabolic compartmentation. In: Lajtha A, editor. Glutamine, glutamate, and GABA in the central nervous system. New York: Springer; 1982. p. 379–389.

3. Bell JK, Yennawar HP, Wright SK, Thompson JR, Viola RE, Banaszak LJ. Structural analyses of a malate dehydrogenase with a variable active site. J Biol Chem. 2001;276:31156–31162.

4. Benesi HA, Hildebrand JH. A spectrophotometric investigation of the interaction of iodine with aromatic hydrocarbons. J Am Chem Soc. 1949;71:2703–2707.

5. Berndsen CE, Bell JK. The structural biology and dynamics of malate dehydrogenases. Essays Biochem. 2024;68:57–72.

6. Bernstein LH, Everse J. Studies on the mechanism of the malate dehydrogenase reaction. J Biol Chem. 1978;253:8701–8707.

7. Bernstein LH, Grisham MB, Cole KD, Everse J. Substrate inhibition of the mitochondrial and cytoplasmic malate dehydrogenases. J Biol Chem. 1978;253:8697–8701.

8. Braidy N, Guillemin GJ, Mansour H, Chan-Ling T, Poljak A, Grant R. Age related changes in NAD+ metabolism oxidative stress and Sirt1 activity in Wistar rats. PLoS One. 2011;6:e19194.

9. Breiter DR, Resnik E, Banaszak LJ. Engineering the quaternary structure of an enzyme: construction and analysis of a monomeric form of malate dehydrogenase from Escherichia coli. Protein Sci. 1994;3:2023–2032.

10. Chen WW, Freinkman E, Wang T, Birsoy K, Sabatini DM. Quantification of matrix metabolites reveals the dynamics of mitochondrial metabolism. Cell. 2016;167:162–177.

11. Classen S, Hura GL, Holton JM, et al. Implementation and performance of SIBYLS: a dual endstation small-angle X-ray scattering and macromolecular crystallography beamline. J Appl Crystallogr. 2013;46:1–13.

12. De Lorenzo L, Stack TMM, Fox KM, Walstrom KM. Catalytic mechanism and kinetics of malate dehydrogenase. Essays Biochem. 2024;68:73–82.

13. Fahien LA, Kmiotek EH, MacDonald MJ, Fibich B, Mandic M. Regulation of malate dehydrogenase activity by glutamate, citrate, alpha-ketoglutarate, and multienzyme interaction. J Biol Chem. 1988;263:10687–10697.

14. Genda T, Nakamatsu T, Ozaki H. Purification and characterization of malate dehydrogenase from Corynebacterium glutamicum. J Biosci Bioeng. 2003;95:562–566.

15. Goward CR, Nicholls DJ. Malate dehydrogenase: a model for structure, evolution, and catalysis. Protein Sci. 1994;3:1883–1888.

16. Grant BJ, Rodrigues APC, ElSawy KM, McCammon JA, Caves LSD. Bio3d: an R package for the comparative analysis of protein structures. Bioinformatics. 2006;22:2695–2696.

17. Hall BA, Kaye SL, Pang A, Perera R, Biggin PC. Characterization of protein conformational states by normal-mode frequencies. J Am Chem Soc. 2007;129:11394–11401.

18. Hopkins JB. BioXTAS RAW 2: new developments for a free open-source program for small-angle scattering data reduction and analysis. J Appl Crystallogr. 2024;57:194–208.

19. Hung YP, Albeck JG, Tantama M, Yellen G. Imaging cytosolic NADH/NAD+ redox state with a genetically encoded fluorescent sensor. PLoS One. 2009;4:e7195.

20. Kikhney AG, Borges CR, Molodenskiy DS, Jeffries CM, Svergun DI. SASBDB: towards an automatically curated and validated repository for biological scattering data. Protein Sci. 2019;29:66–75.

21. Kim G, Lee S, Levy Karin E, et al. Easy and accurate protein structure prediction using ColabFold. Nat Protoc. 2024.

22. Martinez-Vaz BM, Howard AL, Jamburuthugoda VK, Callahan KP. Insights into the regulation of malate dehydrogenases: inhibitors, activators, and allosteric modulation. Essays Biochem. 2024;68:173–181.

23. McCue WM, Finzel BC. Structural characterization of human cytosolic malate dehydrogenase I. ACS Omega. 2022;7:207–214.

24. McKenna MC. The glutamate-glutamine cycle is not stoichiometric. J Neurosci Res. 2007;85:3347–3358.

25. Mullinax TR, Mock JN, McEvily AJ, Harrison JH. Regulation of mitochondrial malate dehydrogenase. J Biol Chem. 1982;257:13233–13239.

26. Olsson MHM, Søndergaard CR, Rostkowski M, Jensen JH. PROPKA3. J Chem Theory Comput. 2011;7:525–537.

27. Pelikan M, Hura GL, Hammel M. Structure and flexibility within proteins as identified through SAXS. Gen Physiol Biophys. 2009;28:174–189.

28. Pettersen EF, Goddard TD, Huang CC, et al. UCSF ChimeraX. Protein Sci. 2021;30:70–82.

29. Rosenberg DJ, Hura GL, Hammel M. Size exclusion chromatography coupled SAXS. Methods Enzymol. 2022;677:191–219.

30. Schneidman-Duhovny D, Hammel M, Tainer JA, Sali A. FoXS and related tools. Nucleic Acids Res. 2016;44:W424–W429.

